# Improving the robustness of engineered bacteria to nutrient stress using programmed proteolysis

**DOI:** 10.1101/2021.10.04.463040

**Authors:** Klara Szydlo, Zoya Ignatova, Thomas E. Gorochowski

## Abstract

The use of short peptide tags in synthetic genetic circuits allows for the tuning of gene expression dynamics and release of amino acid resources through targeted protein degradation. Here, we use elements of the *Escherichia coli* and *Mesoplasma florum* transfer-messenger RNA (tmRNA) ribosome rescue systems to compare endogenous and foreign proteolysis systems in *E. coli*. We characterize the performance and burden of each and show that while both greatly shorten the half-life of a tagged protein, the endogenous system is approximately ten times more efficient. Based on these results, we then demonstrate using mathematical modelling and experiments how proteolysis can improve cellular robustness through targeted degradation of a reporter protein in auxotrophic strains, providing a limited secondary source of essential amino acids that help partially restore growth when nutrients become scarce. These findings provide avenues for controlling the functional lifetime of engineered cells once deployed and increasing their tolerance to fluctuations in nutrient availability.

## Introduction

Prokaryotic protein degradation is an essential cellular quality control mechanism and plays a crucial role in eliminating damaged and/or non-functional proteins ^1–3^. It is enabled by a network of ATP-dependent proteases and adaptors that recognize specific motifs in misfolded proteins, or degrons ^4,5^. Protein degradation in bacteria is mediated by the prokaryotic transfer-messenger RNA (tmRNA) ribosome rescue system, where an SsrA peptide tag is added C-terminally to nascent polypeptides, targeting them for degradation by several endogenous proteases ^6^. These include ClpXP, ClpAP, FtsH and Lon, with ClpXP and ClpAP being the most active in *Escherichia coli*, degrading over 90% of SsrA-tagged proteins ^1,3,7^. The tagging of proteins for degradation has gained interest in the field of synthetic biology as it allows for specific and controllable protein degradation and has been used to modulate protein turnover rates, investigate protein function by reducing intracellular concentrations, and for the tuning of dynamic processes (e.g., modulating the period of genetic oscillators) ^8–11^.

The SsrA peptide-tag system is conserved across prokaryotic species, but the tags vary in their amino acid composition and length ^8,12–14^. The *E. coli* SsrA tag is the most extensively characterized, and its last three amino acids, ‘LAA’, determine the tag strength and the rate of tagged protein degradation ^8^. Variants of these critical residues such as ‘LVA’, ‘AAV’ and ‘ASV’ result in different degradation rates, with ‘LAA’ and ‘LVA’ rendering tagged-GFP more unstable than the ‘AAV’ or ‘ASV’ variants ^8^. The growing knowledge of *E. coli* proteases and their dependency on auxiliary adaptor proteins has also allowed for controllable modulation of protein half-lives and degradation ^2,15,16^. For example, the degradation of proteins tagged with an *E. coli* tag variant ‘DAS’ is mediated by the induction of the SspB adaptor protein in *Bacillus subtilis* ^14^.

Using SsrA tags from distinct species offers another level of control over protein degradation. The simultaneous use of multiple tags in parallel supports the construction of more complex systems where degradation of multiple proteins can be independently controlled. Several SsrA tags from other species have been characterized ^13,14,17^, including that of *Mesoplasma florum* ^12^. This is targeted by the efficient *M. florum* Lon protease that acts orthogonally to the endogenous *E. coli* system, making it possible to use both systems simultaneously in *E. coli* cells ^12^. Previous studies have identified regions of the *M. florum* tag which are crucial for recognition by *E. coli* and *M. florum* proteases, leading to the development of variants of the *M. florum* tag through the deletion of non-essential regions or replacement of residues with other amino acids ^10,18^. Furthermore, the specificity of the endogenous *M. florum* Lon protease to the cognate *M. florum* SsrA tag has enabled the development of inducible orthogonal protein degradation systems in *E. coli* with diverse applications, including the ability to control the behavior of synthetic circuits such as toggle switches ^10–12,18^.

Whilst targeted protein degradation has seen widespread use in tuning the function of genetic parts and circuits, much less attention has been placed on its use in its more native context. Specifically, using protein degradation to help recycle essential amino acid resources when nutrient stress occurs^19,20^. Although such capabilities are less important when cells are grown in the rich and carefully controlled conditions of the lab, when deploying an engineered system into real world environments like your gut or the soil, high variability in nutrient availability is inevitable and cells must be able to react ^21–24^. Therefore, having programmable systems to help buffer cells from these effects is important and warrants further investigation.

Here, we attempt to address this need by exploring how endogenous and heterologous protein degradation systems can be used to manage reservoirs of amino acids that are locked up in stable non-endogenous proteins that can then be subsequently released when needed. We explore the suitability of endogenous and heterologous proteolysis systems for implementing this type of system and show using auxotrophic strains how targeted release of amino acids from a reporter protein enables the partial recovery of growth when an essential amino acid becomes scarce in the growth media. Our proof-of-concept systems offer inspiration for developing new cellular chassis that are more robust to nutrient fluctuations, as well as opening avenues to constrain the functional “shelf-life” of a cell by providing an internal amino acid reservoir with a limited capacity, acting somewhat like a biological battery.

## Results

### Assessing the proteolytic activities of E. coli and M. florum SsrA tags

To gain an insight into the effectiveness of different proteolytic tags, we compared the activities of the *E. coli* and *M. florum* proteolysis systems by assembling genetic constructs where an *eGFP* (GFP) reporter gene was tagged with one of two proteolysis tags. Specifically, we used the *E. coli* (Ec; AANDENYALAA) and *M. florum* (Mf; AANKNEENTNEVPTFMLNAGQANYAFA) SsrA tag sequences which were codon optimized for expression in *E. coli* (**Materials and Methods**) and fused these to the C-terminus of GFP whose expression was under the control of an isopropyl β-D-1-thiogalactopyranoside (IPTG) inducible promoter (P_*lac*_). In this way, GFP was synthesized bearing one of two peptide tags, targeting it for proteolytic degradation by each of our chosen systems (**Figure 1A**). Because the Mf tag is specifically recognized by its cognate Lon protease from *M. florum* (Mf-Lon), which is not present in *E. coli*, we also constructed a separate plasmid where a codon-optimized *lon* gene from *M. florum* ^12^ was expressed under the control of an arabinose-inducible promoter (P_*BAD*_).

**Figure 1:**
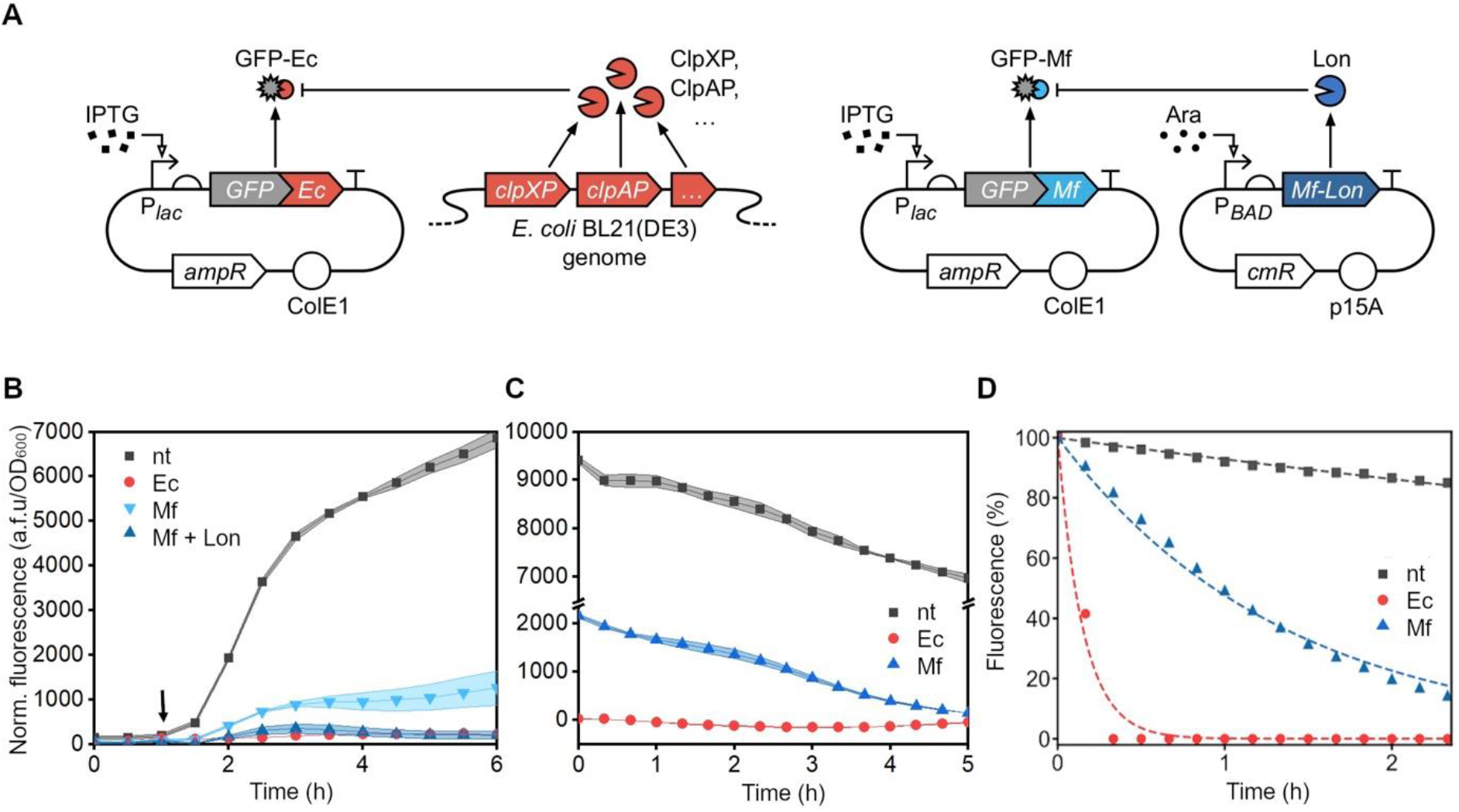
*E. coli* and *M. florum* proteolysis systems used for targeted protein degradation in *E. coli*. (**A**) Schematic of the proteolysis systems. GFP is expressed with *E. coli* (Ec) or *M. florum* (Mf) SsrA tags, which mark it for degradation by endogenous proteases, or the orthogonal plasmid-borne Mf-Lon protease, respectively. (**B**) GFP fluorescence normalized to cell density of *E. coli* BL21(DE3) cells expressing non-tagged GFP (nt), GFP-Ec (Ec) or GFP-Mf without and with the co-expression of Mf-Lon (Mf and Mf + Lon, respectively). Arrow indicates timepoint of GFP induction. (**C**) GFP fluorescence normalized to cell density of cells expressing untagged GFP (nt), GFP-Ec (Ec), or GFP-Mf (Mf) after removal of inducer, whilst maintaining Mf-Lon expression in the case of GFP-Mf. (**D**) % Fluorescence normalized to the time of removal of the inducer of cells expressing untagged GFP (nt), GFP-Ec (Ec), or GFP-Mf (Mf). Curves are fitted to first order exponential decay. Data are means ± SD (*n* = 3 independent biological replicates).

To assess the performance of the two tags, we expressed untagged GFP, GFP-Ec, or GFP-Mf alone and simultaneously with Mf-Lon in *E. coli* BL21(DE3) cells and measured cell growth and fluorescence (**Figure S1**). We observed almost no fluorescence in cells expressing GFP-Ec compared to cells expressing untagged GFP (2.8% at 6 h), indicating that the *E. coli* tag was effective in targeting the tagged protein for degradation by endogenous proteases (**Figure 1B**). In contrast, GFP-Mf when expressed alone, saw reduced, though nevertheless substantial levels of GFP, suggesting that most, but not all, of this protein escaped the endogenous *E. coli* proteases (**Figure 1B**). This was confirmed with additional experiments where the Mf-Lon expressing plasmid was both absent and present, corroborating previous findings ^10,18^ (**Figure S2**). As expected, further induction of Mf-Lon protease caused a 76% drop in GFP-Mf fluorescence, supporting the notion that the Mf tag is specifically recognized (**Figure 1B**). The fluorescence observed from cells expressing the untagged GFP remained largely the same upon induction of the Mf-Lon protease, demonstrating the specificity of the protease for the Mf tag (**Figure S3**).

To further compare the efficiency of the Ec and Mf tags, we induced the expression of untagged GFP, GFP-Mf, or GFP-Ec and after 5 hours removed the inducer. After allowing for GFP maturation ^25^, we then monitored the degradation rate of each GFP variant by the drop in fluorescence and calculated their half-lives (**Figure 1C,D**; **Materials and Methods**). The fluorescence levels of cells containing GFP-Ec remained low throughout, indicating that even strong expression rates could not overcome the endogenous protein degradation. From this data, we found GFP, GFP-Ec and GFP-Mf to have half-lives of 565 min, 6 min, and 56 min, respectively. These numbers support the high efficiency of the endogenous *E. coli* proteases with half-lives being almost ten times shorter than when using the *M. florum* system. However, the Mf-tag did still cause an increased turnover rate, with GFP-Mf exhibiting a half-life less than a tenth of the untagged GFP.

To assess how general these results were, we further tested if each system functioned similarly in the industrially relevant *E. coli* BL21 (DE3) star strain, in which RNase E has been knocked out for higher mRNA stability (**Figure S4**). As expected, we observed longer half-lives for each tagged GFP of 14 min and 424 min for GFP-Ec and GFP-Mf, respectively, and virtually no measurable degradation of the untagged GFP, which was likely due to the increased stability of mRNA in this strain **(Figure S4**).

We also explored the impact of amino acid recycling when a cell is further burdened by a large genetic regulatory circuit. We chose to use a large 3 input, 1 output genetic logic circuit called 0xF6 designed by the Cello software ^26^ and composed of 9 transcription factors. Our existing strains containing the untagged GFP and GFP-Ec were co-transformed with the pAN3938 plasmid encoding this circuit and an assessment of growth carried out (**Figure S5**). As expected, growth was significantly affected by the addition of the 0xF6 genetic circuit. Following an exponential growth which was undistinguishable from that of GFP-nt expressing cells, the GFP-Ec expressing cells gave way to a much slower growth phase, presumably once the circuit began functioning at its full potential and reaching high levels of expression. Despite not seeing any effect of the Ec tag in the exponential growth phase of the cells, we observed that in the second slower growth phase, cells expressing GFP-Ec grew to a higher density than cells expressing GFP-nt. This indicates that, once the expression of the 0xF6 circuit had started to affect the cells, the expression of tagged GFP was able to provide some relief from this burden.

### Dynamic and targeted control of protein degradation using the M. florum SsrA system

A potential advantage of using the *M. florum SsrA* tag system in *E. coli* for the recycling of amino acids is the ability to dynamically control its expression to coincide with an increased demand for resources (e.g., during starvation conditions). This reduces the strength at which tagged proteins acting as a reservoir of amino acids need to be expressed as their turn-over rate can be kept low to ensure long-term protein stability when recycling is not required. Such a method is not possible with the endogenous system as it is continually active and needed by the host cell. Therefore, stronger, and continual expression of the tagged protein is necessary to maintain a similar sized pool of reserve protein.

We carried out several time-course experiments to investigate the precise dynamics of the Mf-tag system in this context where GFP-Mf expression was induced at *t* = 0 and Mf-Lon simultaneously induced or induced 1 or 2 hours after GFP-Mf induction (**Figure S6A**). We found that only simultaneously inducing Mf-Lon with GFP-Mf resulted in increased degradation of GFP-Mf, while sequential induction of Mf-Lon after 1 or 2 h caused barely noticeable drops in fluorescence (4% and 3%, respectively).

This result was unexpected given that Mf-Lon has been shown to function efficiently in *E. coli* ^10,18^, but is likely due to the varying expression strengths of the GFP-Mf reporter and Mf-Lon protease, which reside on different plasmids and which are driven by different promoters (**Figure 1**). To test this theory, we carried out additional experiments where Mf-Lon expression was induced 2 hours before induction of GFP-Mf to allow further time for its accumulation (**Figure S6A**). We found that the initial increase in fluorescence when Mf-Lon was induced simultaneously with GFP-Mf, was negated when Mf-Lon was induced 2 hours prior, suggesting that expression and maturation of Mf-Lon occurs quickly, and efficient GFP degradation could occur. Nevertheless, the rate of fluorescence increase from 3 hours after GFP-Mf induction was almost identical (**Figure S6B**), indicating that the concentration of Mf-Lon achieved when expressed from a P_*BAD*_ promoter and medium-copy plasmid (p15A origin; ∼10 copies per cell) is unable to significantly reduce GFP-Mf levels.

### Recovering cell growth by amino acid recycling

A major challenge when developing genetic circuits is managing the burden they place on shared cellular resources ^27–32^. The expression of a genetic construct will sequester key cellular machinery like ribosomes and may exhaust amino acid supplies, which in turn can impact overall cell physiology and protein synthesis ^33–36^, alter translation dynamics^28^ and trigger stress responses^37,38^. A reason for this large impact is that circuit components are often strongly expressed and designed to be highly stable, causing a large portion of the cell’s resources to become locked away from use in endogenous processes. Several studies have engineered ways to mitigate the burden placed on the cell; limiting recombinant protein expression via negative feedback loops, or reducing translational demand by splitting recombinant protein synthesis between endogenous and orthogonal ribosomes^30,37,39^. However, it has also been observed that supplementing the growth media of cells expressing recombinant proteins with amino acids can enhance growth rate and protein production ^38^. Consequently, we hypothesized that by increasing amino acid turnover of heterologous protein products through targeted proteolysis, we would be able to help mitigate the burden a genetic circuit places on its host cell.

To test this idea, we measured the growth rate of cells expressing tagged and untagged GFP under the control of the same strong P_*lac*_ promoter (**Figure 1A**). We reasoned that the expression of the tagged GFP would place less of a burden on the host compared to the untagged version, due to increased recycling of amino acids ^27,38,40^. Although the expression of any GFP protein will reduce cell growth rate, the reduction in growth rate for the first three hours of induction was smaller for cells expressing GFP-Ec (41%) and GFP-Mf with Mf-Lon (37%), compared to cells expressing untagged GFP (51%) (**Figure S7**). This suggests that while the expression of a recombinant protein will always cause a burden, this burden is partially alleviated by more effective recycling of these products, making these resources accessible to endogenous processes.

It is known that protein degradation is elevated under various stress conditions, possibly as a way to increase the availability of amino acids for synthesis of stress-related proteins ^19,41^. Furthermore, as part of the *E. coli* stringent response to nutrient limitations, there is an increase in the level of amino acid biosynthesis enzymes, to meet the demand for amino acids ^42^. Considering this, we asked whether the potential benefit of using tagged proteins might increase when the host cell experienced nutrient related stress. We reasoned that increased recycling of a heterologous pool of proteins could benefit a host cell where nutrients to synthesize amino acids had become scarce in the environment.

To assess the feasibility of this approach, we developed a simple mathematical model to capture the key flows of a hypothetical essential resource in the cell (e.g., an amino acid the cell is unable to synthesize) and its impact on cell growth (**Figure 2A**). The model consisted of three ordinary differential equations that track the concentrations of a shared resource that is either available for use within the cell (*N*_*c*_), is actively in use by endogenous proteins (*P*_*e*_), or is locked up in foreign heterologous proteins (*P*_*f*_):

**Figure 2:**
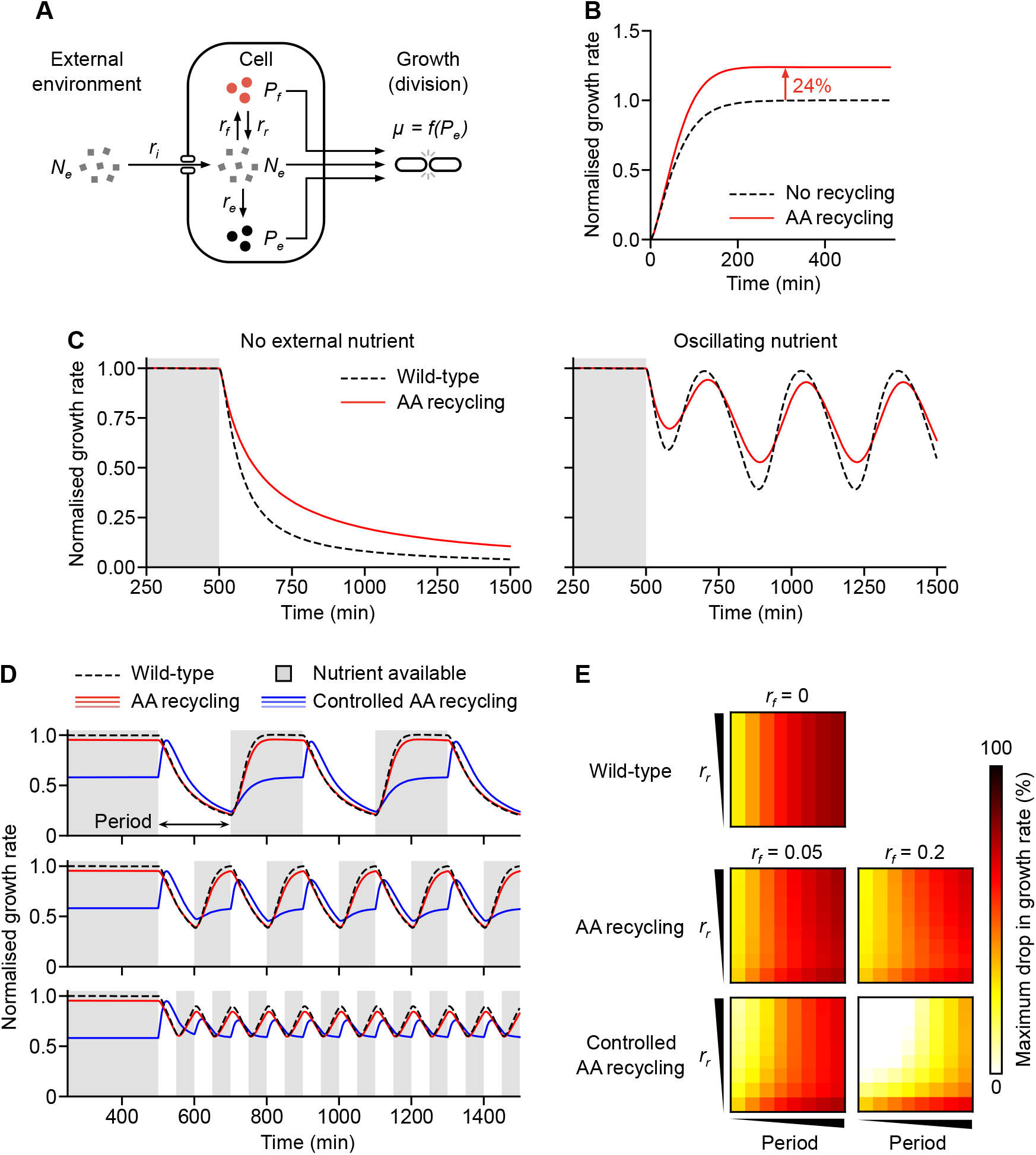
Model capturing the benefits of amino acid recycling. (**A**) Schematic of the model. *N*_*e*_ denotes the external resource concentration, *N*_*c*_, *P*_*e*_ and *P*_*f*_ denote the concentration of a key resource (i.e., an amino acid that cannot be natively produced), available within the cell, locked up in endogenous or in heterologous proteins, respectively. *r*_*i*_ denotes the cellular import rate of resources, which can be divided into *r*_*e*_, the rate at which resources are converted into endogenous proteins, and *r*_*f*_, the rate at which resources are converted into foreign heterologous proteins. *μ* = *f*(*P*_*e*_) captures cell growth and dilution of resources by cell division. (**B**) Simulation of the normalized cell growth rate in a strain expressing recombinant proteins (i.e., with no amino acid recycling), and in a strain expressing tagged heterologous proteins (i.e., with amino acid recycling). (**C**) Simulation of the effects of nutrient stress on normalized cell growth rate expressing tagged proteins (i.e., with continual amino acid recycling; ‘AA recycling’), and in a strain expressing no heterologous protein (‘wild-type’). The external resource (nutrient) is continually present for the first 500 min (grey shaded region), then either removed completely, or oscillating nutrient levels are applied after this time. In all cases, growth rate is normalized to the steady state growth rate when the external resource is present (i.e., *N*_*e*_ = 1). (**D**) Time-series of normalized growth rate (to wild-type cells when the external resource is present) of cells exposed to a cycling of external nutrient absence (white regions) and presence (grey shaded regions). Responses shown for wild-type cells (black dashed line), cells with continual amino acid recycling (red lines; *r*_*f*_ = 0.1 protein resource^−1^ min^−1^, light–dark: *r*_*r*_ = 0.1, 0.2, 0.4 resource protein^−1^ min^−1^), and cells where amino acid recycling is only active when external nutrient is absent (blue lines; *r*_*f*_ = 0.1 protein resource^−1^ min^−1^, light–dark: *r*_*r*_ = 0.1, 0.2, 0.4 resource protein^−1^ min^−1^). Panels from top to bottom show varying length of period that nutrient is absent from the external environment. (**E**) Heat maps showing how varying heterologous protein production (*r*_*f*_ = 0, 0.05, 0.2 protein resource^−1^ min^− 1^) and recycling (*r*_*r*_ = 0.001, 0.015, 0.029, 0.043, 0.057, 0.072, 0.086, 0.1 resource protein^−1^min^−1^) rates, and period (25, 50, 75, 100, 125, 150, 175, 200 min) of the nutrient/resource availability cycling (see panel D) affect the maximum percentage drop in the initial steady state growth rate when external nutrient is present.

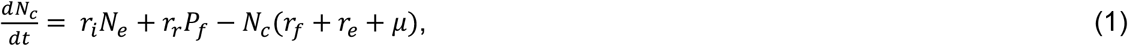

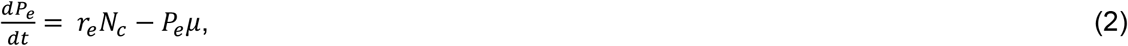

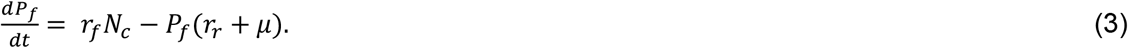

Here, *N*_*e*_ is the external resource concentration outside the cell with a cellular import rate of *r*_*i*_, *r*_e_ and *r*_*f*_ are the rates that available resources within the cell are converted into endogenous or heterologous proteins, respectively, and *r*_*r*_ is the recycling rate of the heterologous proteins (e.g., due to targeted proteolysis). Cellular growth and the associated dilution (by cell division) of all resources was captured by *μ* = 0.1*P*_*e*_. Parameters were chosen such that overall growth rate of the cell was consistent with *E. coli* data (i.e., having a division time ∼25 min) and that relative internal transport, production and degradation rates were biologically realistic (**Materials and Methods**).

Using this model, we simulated cells expressing tagged and untagged proteins (**Figure 2B**) and exposed these cells to several external environmental shifts to temporally vary the resources available (**Figure 2C**). In the first shift, we removed all resources from the environment at 500 min, and in the second, at the same time point, we applied an oscillating external nutrient concentration. In both cases, we compared cells not producing any heterologous protein (i.e., *r*_*f*_ = 0) to those producing a recombinant protein that is subsequently recycled for reuse by the cell. We then measured their response in terms of growth rate normalized to when the external nutrient was continually present (i.e., the steady state growth rate when *N*_*e*_ = 1). In both cases, the model showed a reduction in the relative impact on growth rate to changes in environmental availability (**Figure 2C**), demonstrating the ability for a recycled internal reservoir of a heterologous resource to act as a reserve that can help buffer the cell temporarily from environmental change. It should be noted that inclusion of a heterologous resource pool and its recycling does have an impact on cellular growth rate. However, for some applications (e.g., excitable systems that are sensitive to even minor fluctuations in cellular behaviors ^43^), it may be preferable to have a more consistent performance when faced with environmental variability.

Even though our previous experiments had shown a limited capacity to dynamically vary protein degradation rates, we also explored how future controllable amino acid recycling systems might compare to a simpler system where heterologous proteins are continually recycled. We ran simulations of our model where no heterologous protein pool was present (‘Wild-type’; *r*_*f*_ = 0) and where a heterologous protein pool was continually recycled (‘AA recycling’) or only recycled upon removal of nutrient from the environment (‘Controlled AA recycling’). We allowed each system to reach an initial steady state with the external nutrient present (i.e., *N*_*e*_ = 1 until *t* = 500 min), then provided alternating time periods where the nutrient was completely removed from the environment (i.e., *N*_*e*_ = 0) and then made available again. We tracked the varying growth rate over time to assess the impact on the cells (**Figure 2D**).

As expected, wild-type cells displayed large drops in growth rate upon removal of external nutrient that was proportional to the length of the removal period and fast recovery was seen upon reintroduction of the external nutrient. Activation of constant recycling saw a minor reduction in growth rate compared to the wild-type cells. However, this allowed for the drop in growth rate upon nutrient removal to be a smaller fraction of the initial growth rate. In contrast, controlled amino acid recycling that was active only when the essential nutrient was removed from the environment showed two different features. First, because recycling was only active upon removal of the external nutrient, for normal conditions there was no recycling of the heterologous protein and so a larger impact was seen on the normal growth rate compared to when continual recycling was used (e.g., lower initial normalized growth rates for blue lines in **Figure 2D**). Second, removal of the external nutrient led to a transient increase in growth rate as recycling was activated. However, as the pool of heterologous protein was consumed the growth rate also then began to drop. If the period of nutrient switching was short enough though, it was possible for no reduction below the initial growth rate to be seen (e.g., blue lines in bottom panel of **Figure 2D**).

We also generated heat maps showing the percentage drops in growth rate from an initial steady-state growth rate where the essential nutrient was abundant in the environment (i.e., *N*_*e*_ = 1) for varying recycling rates (*r*_*r*_) and periods of removal from the environment, across both low and high rates of heterologous protein production (*r*_*f*_). These simulations revealed that as expected, when no heterologous protein production is present, growth rate drops with the length of time (period) the essential nutrient is removed from the environment (**Figure 2E**). The expression and continual recycling of a heterologous protein was able to reduce these drops in growth rate, and this effect was enhanced if the internal pool of heterologous protein was sufficiently large and recycled at a sufficiently high rate (i.e., high *r*_*r*_ and *r*_*f*_). For the controlled amino acid recycling, we found that for particular combinations of heterologous protein production and recycling rates and shorter periods of nutrient removal, drops in growth rate could be completely eradicated (white regions in **Figure 2E**). This suggests that controlled amino acid recycling (if sufficiently rapid) is a feasible strategy for completely shielding a cell from environmental nutrient fluctuations. However, trade-offs in the size of the internal heterologous protein pool and the rate of recycling affect the ability to robustly respond to differing lengths of nutrient fluctuation.

### Buffering auxotrophic cells from environmental amino acid fluctuations

To test some of the model predictions, we used auxotrophic *E. coli* strains RF10 ^44^ (Δ*lysA*) and ML17 ^45^ (Δ*glnA*), which are unable to synthesize lysine and glutamine, respectively. This allowed us to tightly control endogenous amino acid levels by modulating the external supply in the media. Furthermore, lysine and glutamine are amongst the most abundant amino acids in our GPF reporter (8.4% and 6.7% of the total amino acid composition, respectively) offering suitable reservoirs of these key resources. We initially tested the ability of the endogenous Ec tag system to enhance cell growth as we had previously found that it resulted in faster degradation of GFP compared to the orthogonal Mf tag system. We grew each of the strains expressing untagged GFP and GFP-Ec in nutrient-rich media to allow for a buildup of the recombinant protein. Following this, cells were switched to minimal media, effectively removing the source of all external amino acids, and for our auxotrophic strains, completely removing access to lysine and glutamine, respectively.

Consistent with our model predictions, we found that both strains expressing GFP-Ec exhibited a higher growth rate than cells expressing untagged GFP; 0.128 and 0.15 h^−1^ for GFP-Ec versus 0.08 and 0.091 h^−1^ for GFP-nt for the Δ*lysA* and Δ*glnA* strains, respectively. This equated to an increase in growth rate of 60% and 65% for the Δ*lysA* and Δ*glnA* strains, respectively (**Figure 3A-B**). We suspect the higher growth rates are due to the degradation of GFP-Ec, which is supported by the lower fluorescence levels (**Figure 3C**). Addition of lysine or glutamine (7 mM) to the medium for the respective untagged GFP-expressing auxotrophic strains saw a marked increase in cell growth rate from 0.008 to 0.13 h^−1^ for the Δ*lysA* strain when lysine was present, and from 0.091 to 0.193 h^−1^ for the Δ*glnA* strain when glutamine was present (**Figure 3B**). This indicated that glutamine and lysine were the major limiting factors for cell growth and that recycling of the internal heterologous protein reservoir was able to partially buffer this impact (37% and 31% recovery for Δ*lysA* and Δ*glnA*, respectively).

**Figure 3:**
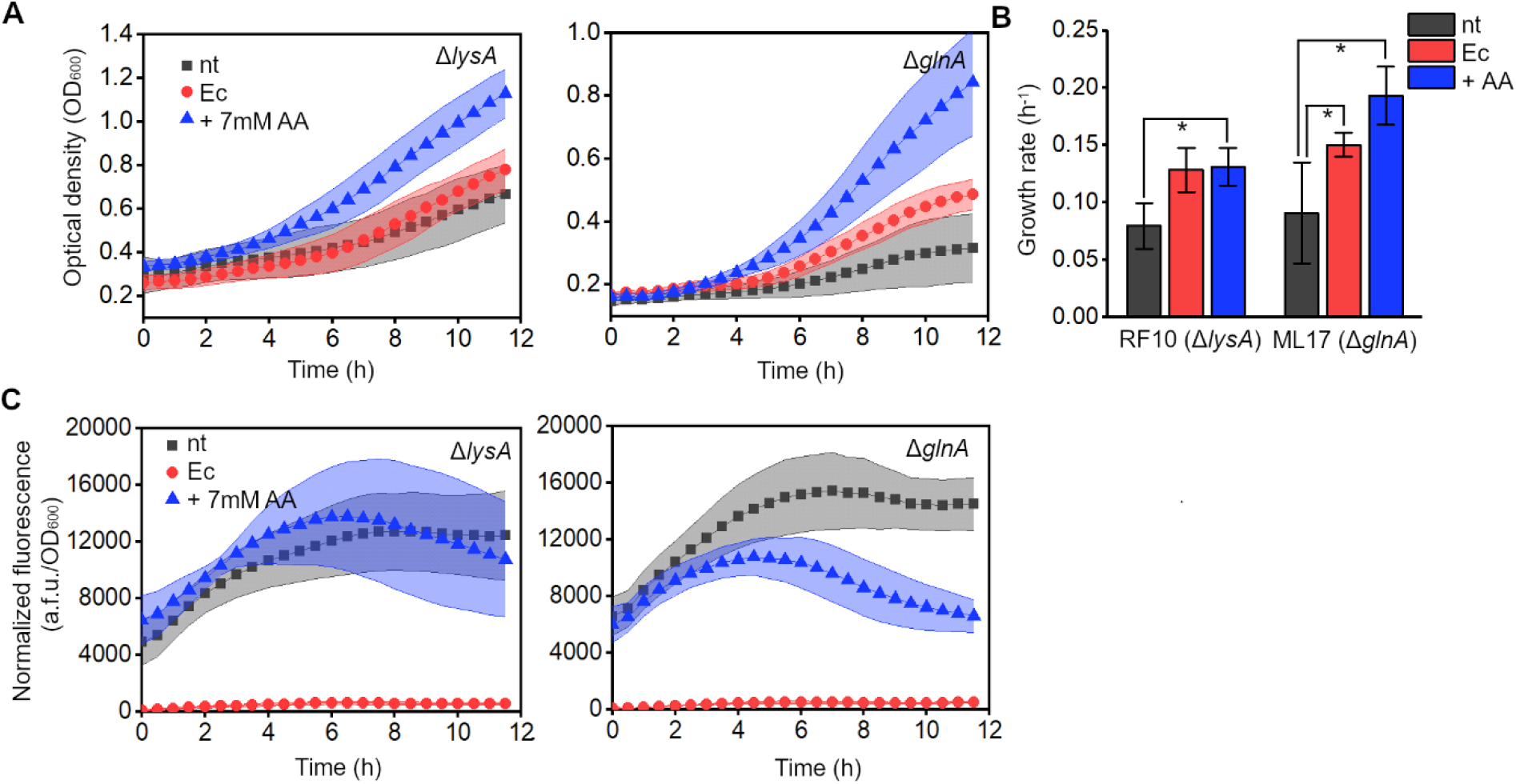
Targeted GFP degradation provides amino acids to auxotrophic strains upon nutrient limitation. (**A**) Growth of the RF10 (Δ*lysA*) and ML17 (Δ*glnA*) strains in minimal medium, expressing untagged GFP (nt) or GFP-Ec (Ec) with the addition of 7 mM lysine or glutamine supplement. Data are means ± SD (**B**) Quantification of the exponential growth rates of cells. (\**p* < 0.05, as compared to nt condition for each strain, with 2-sample *t*-test). Data are means ± SE (**C**) GFP fluorescence normalized to cell density of the RF10 (Δ*lysA*) and ML17 (Δ*glnA*) strains, expressing untagged GFP (nt) or GFP-Ec (Ec) with the addition of 7 mM lysine or glutamine supplement. Data are means ± SD (*n* = 5 independent biological replicates).

Having shown that using the native *E. coli* tag could render cells more robust in the face of amino acid limitations, we next investigated whether the same effect could be seen when using the orthogonal Mf tag system. As mentioned previously and shown by our modelling, an orthogonal system would confer benefits over the endogenous one as it could target degradation in a dynamic and controllable manner. Again, we grew strains co-expressing untagged GFP and Mf-Lon, or GFP-Mf and Mf-Lon in rich media, before switching them to minimal media to eliminate external sources of nutrients. We observed a higher growth rate in both strains when expressing GFP-Mf compared to untagged GFP (0.076 and 0.06 h^−1^ for GFP-Mf versus 0.033 and 0.014 h^−1^ for GFP-nt for the Δ*lysA* and Δ*glnA* strains, respectively). The increase in growth of over 2-fold for the Δ*lysA* strain and over 4-fold for the Δ*glnA* strain corroborated our findings from the model and endogenous tag system (**Figure 4A-B**). This again could be attributed to the degradation of GFP-Mf as indicated by the lower fluorescence levels (**Figure 4C**). We also found that addition of 10 mM lysine or glutamine to the medium recovered the growth of cells expressing untagged GFP and Mf-Lon; 0.033 h^−1^ increased to 0.068 h^−1^ in the Δ*lysA* strain, and 0.014 h^−1^ increased to 0.057 h^−1^ in the Δ*glnA* strain (**Figure 4A-B**). In contrast to the previous results, specifically in Δ*glnA*, the amino acid supplementation, even at fairly high concentrations of 10 mM did not result in an increased growth rate compared to cells with already induced GFP degradation, suggesting that maximum cell growth had already been achieved under these conditions.

**Figure 4:**
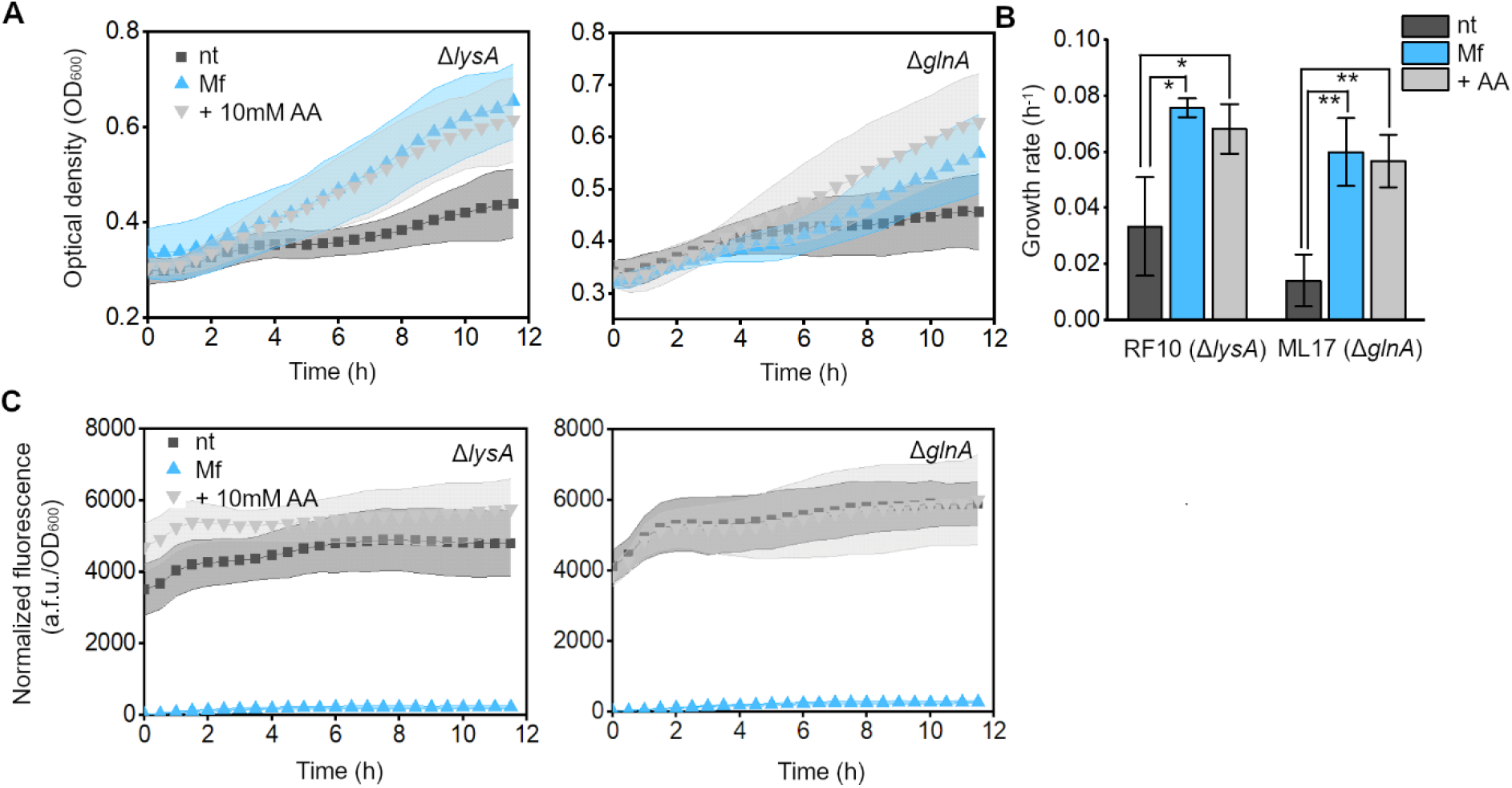
Targeted GFP degradation using the foreign Mf tag system provides amino acids to auxotrophic strains upon nutrient limitation. **(A)** Growth of the RF10 (Δ*lysA*) and ML17 (Δ*glnA*) strains in minimal medium, co-expressing untagged GFP and Mf-Lon (nt), GFP-Mf and Mf-Lon (Mf), or untagged GFP and Mf-Lon (nt) with the addition of 10mM lysine or glutamine supplement (+AA). Data are means ± SD. **(B)** Quantification of the exponential growth rates of cells (\**p* < 0.05, \*\**p* < 0.005, as compared to nt condition for each strain, with 2-sample *t*-test). Data are means ± SE. **(C)** GFP fluorescence normalized to cell density of the RF10 (Δ*lysA*) and ML17 (Δ*glnA*) strains, co-expressing untagged GFP and Mf-Lon (nt), GFP-Mf and Mf-Lon (Mf), or untagged GFP and Mf-Lon (nt) with the addition of 10mM lysine or glutamine supplement (+AA). Data are means ± SD (*n* = 5 independent biological replicates).

We also found that cells expressing both GFP-Mf and Mf-Lon grew faster than cells expressing only GFP-Mf (**Figure 5A-B**). The induction of the Mf-Lon protease enhanced growth of both the Δ*lysA* and Δ*glnA* strains; 0.076 and 0.06 h^−1^, respectively, compared to where GFP-Mf alone was expressed; 0.057 and 0.026 h^−1^, respectively. This suggests that the benefits of increased protein degradation (**Figure 5C**), and therefore a higher level of amino acid recycling, outweigh the cost of expressing two recombinant proteins (the reporter and the protease). Indeed, upon Mf-Lon protease induction, we observed an increase in growth of 33% and 180% for the Δ*lysA* and Δ*glnA* strains, respectively, providing support for increased amino acid recycling within cells enhancing their robustness to nutrient stress. Interestingly, we observed that the Δ*glnA* strain grew slower than the Δ*lysA* strain, and that the benefits of amino acid recycling were more pronounced (**Figures 4 and 5**). This may be due to the fact that glutamine is more common in the *E. coli* proteome than lysine ^46^. Therefore, a lack of endogenous glutamine would have a greater effect on cellular growth when external nutrients were limited and cells were heavily burdened, than a lack of endogenous lysine.

**Figure 5.**
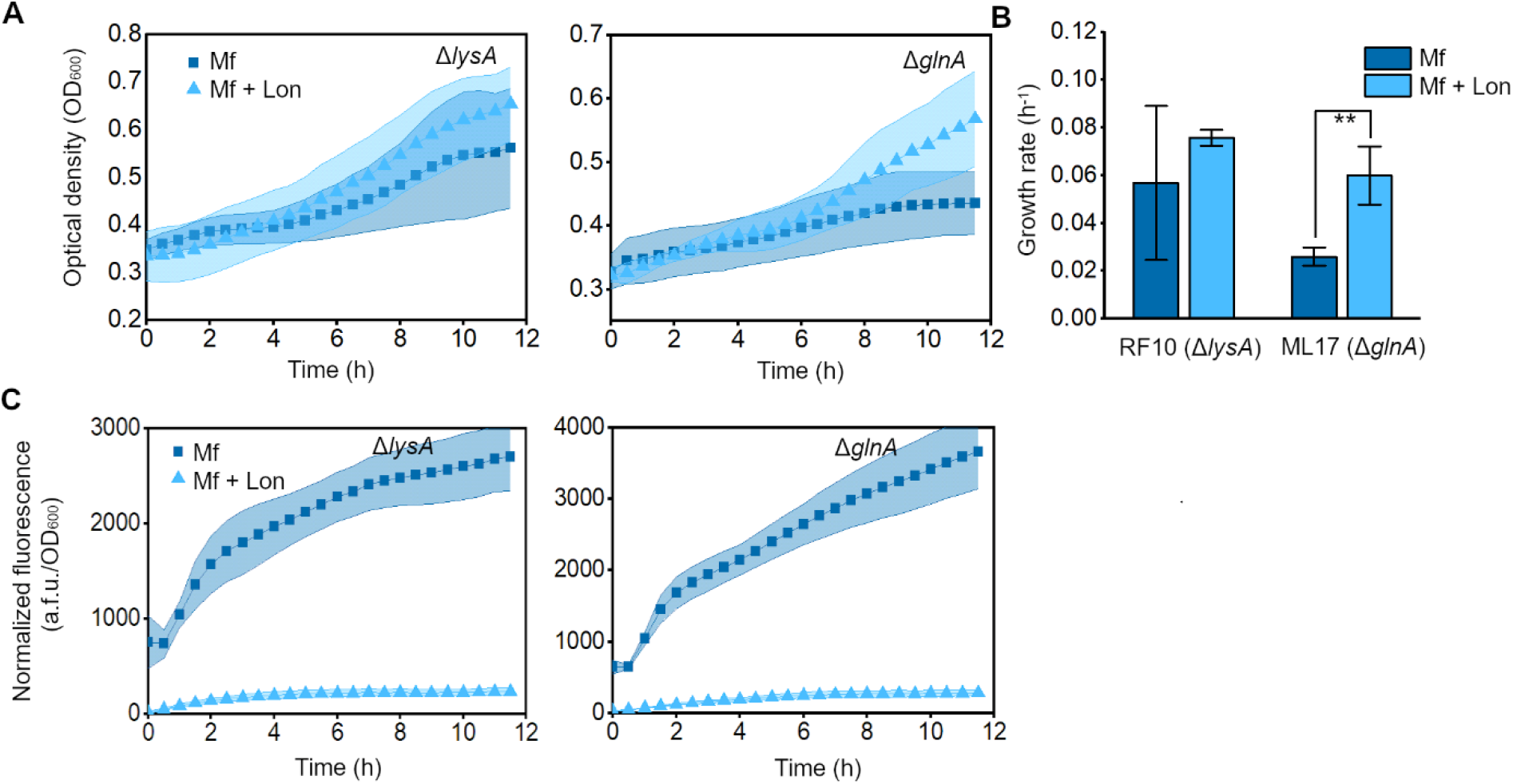
Induction of the orthogonal Mf proteolysis system increases cell robustness against nutrient stress as a result of resource recycling. **(A)** Growth of the RF10 (Δ*lysA*) and ML17 (Δ*glnA*) strains in minimal medium expressing GFP-Mf (Mf), or GFP-Mf and Mf-Lon (Mf + Lon). Data are means ± SD. **(B)** Quantification of the exponential growth rates of cells (\*\**p* < 0.005, as compared to Mf condition, with 2-sample *t*-test). Data are means ± SE. **(C)** GFP fluorescence normalized to cell density of the RF10 (Δ*lysA*) and ML17 (Δ*glnA*) strains expressing GFP-Mf (Mf), or GFP-Mf and Mf-Lon (Mf + Lon). Data are means ± SD (*n* = 5 independent biological replicates).

Together, these results show that targeted degradation of heterologous proteins can be beneficial to cells experiencing severe nutrient stress and be used to buffer growth rate from fluctuations in intracellular levels of amino acids.

## Discussion

In this work, we have directly compared the effectiveness of the endogenous *E. coli* proteolysis system and a similar heterologous system from *M. florum*, and characterized them with the goal of using them as mechanisms for promoting targeted amino acid recycling within *E. coli* cells (**Figure 1**) ^8,12^. We found that the endogenous system was approximately ten times more effective than the *M. florum* system, shortening the half-life of untagged GFP almost 100-fold, and similar results were observed in a second *E. coli* BL21(DE3) star strain, indicating the transferability of this approach. We also observed some crosstalk between these systems, with the reporter protein containing the *M. florum* tag also seeing increased degradation compared to an untagged reporter when the cognate Mf-Lon protease was not present. While characterization of these systems has been performed independently ^8,10,11,18,47^, we believe this study to be the first that directly compares these systems targeting an identical target protein and functioning within the same host cell context.

In addition, we explored the option to activate targeted degradation by externally inducing expression of the *M. florum* system dynamically over time. However, we found that dynamic expression of the Mf-Lon protease in our system was unable to have a significant effect on GFP levels unless the protease was simultaneously induced or induced prior to the target protein (**Figure S6**). This was likely due to the use of different plasmid backbones with different plasmid copy numbers, resulting in strong expression of GFP-Mf that Mf-Lon could not overcome. We also studied the effect of proteolysis tags when the host cell expressed a large genetic circuit, and found that the Ec tag remained effective and supported slow cell growth after the burdensome circuit became established within them (**Figure S5**).

Numerous strategies have been developed to mitigate the burden that genetic circuits and strong heterologous protein expression places on a host cell. Examples include the use of negative regulators, stress feedback sensors, and orthogonal ribosomes ^30,37,39^. In our study, we asked whether the genetic circuit burden could be mitigated by using proteolysis tags to stimulate a higher turnover rate of amino acids, which could be used for the synthesis of endogenous proteins. We found that the reduction in growth rate of cells was indeed smaller when they expressed a tagged protein (**Figure S7**), providing evidence for the benefits of using proteolysis tags when expressing recombinant proteins. Based on these findings, we developed a model that further explored the role that proteolysis could play, specifically under nutrient stress. The model showed that benefits would be amplified when facing external amino acid shortages, especially when activation of proteolysis could be triggered only when necessary (**Figure 2**). Finally, by using auxotrophic strains, we were able to show that recycled heterologous proteins could act as a limited reservoir of amino acid resources, both when using the endogenous and orthogonal tag systems, helping buffer the cell from fluctuations in nutrient availability and partially recover cell growth (**Figures 3 and 4**). We also found that the benefits of constant proteolysis outweighed the costs of expressing two recombinant proteins (**Figure 5**), emphasizing the importance of resource recycling to cells exposed to nutrient stress.

As our ambitions in synthetic biology grow and we begin to consider the construction of entire synthetic cells, understanding how resources flow and are recycled throughout these systems will become crucial. Our demonstration of the benefits of proteolysis tags as a means for amino acid recycling opens new avenues for other approaches controlling nutrient fluxes within a cell and provides a fresh perspective on the use of internal pools of heterologous proteins (or other resources) that can be released when needed to alleviate potential environmental fluctuations. This methodology can be used to help buffer the cell, and the engineered genetic systems they host, from the unavoidable variability that is present within real-world environments and paves the way for creating more reliable and robust biosystems.

## Materials and Methods

### Bacterial strains, media, and cloning

The *E. coli* strain DH5α (*φ80dlacZ ΔM15 Δ(lacZYA-argF)U169 deoR recA1 endA1 hsdR17rK-mK+ phoA supE44 λ– thi-1*) was used for plasmid construction and cloning, and the strains BL21(DE3) (*F – ompT hsdSB (rB– mB–) gal dcm (DE3)*) and BL21(DE3) star (*F – ompT hsdSB (rB– mB–) gal dcm rne131 (DE3)*) used for characterisation of our genetic systems. Cells were grown in Luria-Bertani (LB) media (Roth, X968.4), or minimal media (12.8 g/l Na_2_HPO_4_.7H_2_O, 3 g/l KH_2_PO_4_, 1 g/l NH_4_Cl, 2 mM MgSO_4_, 0.1 mM CaCl_2_, 0.4% glucose). 100 mg/ml ampicillin (Sigma Aldrich, A9393), 50 mg/ml kanamycin (Sigma Aldrich K1377), or 34 mg/ml chloramphenicol (Sigma Aldrich, C0378) were used as selection markers for cloned plasmids. Enhanced green fluorescent protein (eGFP) in the pET16b vector under the IPTG-inducible *Lac* promoter system was C-terminally tagged with the *E. coli* (Ec) tag through site directed mutagenesis: overlap PCR primers were designed which contained the Ec tag sequence. These were phosphorylated and used for PCR with the plasmid backbone. The product was digested with *DpnI* (NEB, R0176S) overnight, and the resulting product PCR purified. A ligation was carried out to circularise the vector, using 10-50 ng of DNA and T4 DNA ligase (Thermo Fisher, EL0011), according to the manufacturer’s instructions. The resulting plasmid was transformed into competent DH5α cells. The *M. florum* tag was codon optimized for expression in *E. coli* by selecting the most highly abundant codons in *E. coli* for the corresponding amino acids (codon sequence: GCT GCA AAC AAG AAC GAG GAA AAC ACC AAC GAA GTA CCG ACC TTC ATG CTG AAC GCA GGC CAG GCT AAC TAT GCA TTC GCA), and GFP was C-terminally tagged with the Mf tag using a digest-and-ligate approach: oligonucleotides were designed to contain the Mf tag sequence, and annealed to create double-stranded DNA fragments, then phosphorylated. The pET16b-eGFP vector was digested with fast digest *BsrgI* and *XhoI* (NEB, R0102S and R0146S) and used in a ligation reaction with the inserts (3:1 ratio) using T4 DNA ligase (Thermo Fisher, EL0011), at 22°C for 4–6 h. Competent DH5α cells were transformed with the resulting product. The *M. florum* Lon protease from the pBAD33 vector (a gift from Robert Sauer; Addgene plasmid 21867) was subcloned into the pSB3C5 plasmid under the *araBAD* promoter using Golden Gate assembly. The primers for the PCR reaction were designed to flank the *Mf-Lon* with *BsmBI* restriction sites and include them into the vector (pSB3C5). The Golden Gate assembly reaction was set up, which included insert:vector in a 4:1 ratio, *BsmBI* (NEB, R0739S), and 1 μl T4 DNA Ligase (Thermo Fisher, EL0011). The following conditions were used for the reaction: 60 cycles of 42°C for 3 min then 16°C for 4 min, followed by 50°C for 5 min and 80°C for 5 min. The resulting product was transformed into *E. coli* DH5α cells. The pAN3938 plasmid encoding the 0xF6 genetic logic circuit ^26^ was a gift from Christopher Voigt, and electrocompetent BL21 cells were co-transformed with either the GFP-Ec or GFP-nt plasmids, and the pAN3938 plasmid.

### Proteolysis tag activity assays

Overnight cultures of BL21(DE3) or BL21(DE3) star cells transformed with pET16b-eGFP-no tag, pET16b-eGFP-Ec, or pET16b-eGFP-Mf and pSB3C5-mfLon were grown for 12-16 h at 37°C 250 rpm, then re-suspended in minimal media with appropriate antibiotics for selection. The cultures were grown to an OD_600_ of 0.4–0.6, then induced with 0.5 mM IPTG (Roth, 2316.3) or 0.2% (w/v) arabinose (Roth, 5118.2). 1% glucose was added to the cultures expressing untagged proteins, to prevent basal expression from the pET16b vector^48^. For degradation assays, cells were pelleted 5 h after induction, washed twice in 1X phosphate-buffered saline (PBS) (137 mM NaCl, 2.7 mM KCl, 10 mM Na_2_HPO_4_, 1.8 mM KH_2_PO_4_) then re-suspended in minimal medium containing the relevant antibiotics, without IPTG. In cultures co-transformed with two plasmids, IPTG induction was stopped, but the second inducer, 0.2% (w/v) arabinose, was added to the medium to induce expression of Mf-Lon. 200 μl of the cultures were grown in a 96-well flat-bottom black plate with clear bottom (Corning, Sigma Aldrich CLS3603-48EA) at 37°C with orbital shaking in a multimode microplate reader (Tecan Spark). Optical density (at 600 nm) and fluorescence measurements (excitation and emission wavelengths of 472 nm and 517 nm, respectively, with a gain of 50) were recorded at discrete intervals. Fluorescence was normalised to the OD_600_ value (a.f.u./OD_600_). Untransformed BL21(DE3) cells were used as a negative control and their normalised autofluorescence values (a.f.u/OD_600_) were subtracted from the normalised fluorescence values (a.f.u./OD_600_) of the cells in different conditions.

### Auxotrophic strain and starvation assays

The RF10 (Δ*lysA*) and ML17 strains (Δ*glnA*) (a gift from Robert Gennis & Toshio Iwasaki Addgene plasmids 62076 and 61912) were transformed with the plasmids developed in this work and grown in LB to an OD_600_ of ∼0.3. For cells transformed with GFP-nt and GFP-Ec, GFP expression was induced with 0.5 mM IPTG for 1 h. For cells transformed with GFP-Mf and Mf-Lon, or GFP-nt and Mf-Lon, GFP expression was induced with 0.5 mM IPTG and Lon expression induced with 0.2% (w/v) arabinose for 1 h. After this, cells were pelleted, washed in 1X PBS, and re-suspended in minimal medium containing appropriate antibiotics for plasmid selection with additional 0.5 mM IPTG to maintain GFP expression, or 0.5 mM IPTG and 0.2% (w/v) arabinose to maintain GFP and Mf-Lon expression. Additionally, 7 or 10 mM of lysine (Sigma Aldrich, L5501) or glutamine (Serva, 22942) were added to positive control samples. The OD_600_ value and fluorescence were then measured as described above using a microplate reader every 10 min over 12 h.

### Data analysis

Python version 3.9.5 and packages matplotlib version 3.3.2, NumPy version 1.19.2, and SciPy version 0.13 were used to fit the degradation data to a first order decay function of the form, *N*(*t*) = 100*e* – λ*t*, where *N*(*t*) is the percentage of remaining fluorescence at time *t* post the start of the degradation curve, and λ the decay constant. The half-live of GFP was then given by *t*_1/2_ = ln(2)/λ. When investigating the dynamics of the Mf-tag system, the rate of GFP production (*F*) was calculated as the gradient of fluorescence values normalized to OD_600_ between 3 and 7 h after induction (**Figure S6**). To obtain values for the growth rate of cells expressing tagged or untagged GFP, the slope of the linear fit to log transformed OD6_00_ values was calculated when cells were in their exponential growth phase, between 5 and 9 h (**Figures 3-5**), or 1.5 and 2.5 h (**Figure S5**) ^49^. It is of note to consider that the cells do not always show standard growth curves with distinct growth phases, due to confounding factors such as the auxotrophic strains used in the experiments, nutrient stress placed on the cells, as well as the expression burden. Therefore, the exponential growth phase of cells is not always distinct. The % drop in growth rate of cells expressing recombinant proteins used growth rate values calculated between 1–4 h growth (Figure S7).To compare whether the growth rates of the auxotrophic strains were statistically significantly different, a 2-sample *t*-test was used. *p* < 0.05 was considered as statistically significant. The statistical analysis was performed, and all plots and slopes of best fit were generated, using OriginLab Pro software (2019 version 64 bit).

### Model parameterization and simulation

Parameters for the model of resource allocation and use were selected based on the assumption that an external resource concentration of *N*_*e*_ = 1 would lead to a realistic cell doubling time (∼25 min) and that internal cellular rates would have biologically feasible relative values. This resulted in simulations with heterologous protein recycling present being simulated with parameters: *r*_*i*_ = 0.015, *r*_*e*_ = 0.02, *r*_*f*_ = 0.2, *r*_*r*_ = 0.01, and with *μ* = 0.1 × *P*_*e*_. In all simulations, initial conditions for all states were set to 0, and the dynamics simulated for 500 min with *N*_*e*_ = 1.0 for the system to reach a steady state before any environmental fluctuations occurred. The model was simulated using Python version 3.9 and the SciPy version 0.13. The code for all simulations is available as **Supplementary Data 1**.

## Supporting Information

The effects of Ec and Mf proteolysis tags on cell fluorescence and growth (**Figure S1**); Off-target degradation by *E. coli* proteases on the Mf tag (**Figure S2**). The specificity of the Lon protease for the Mf tag (**Figure S3**). *E. coli* and *M. florum* proteolysis systems are effective among different *E. coli* strains (**Figure S4**). Programmed proteolysis allows for better cell growth when hosting burdensome genetic circuits (**Figure S5**). The dynamics of the *M. florum* proteolysis system (**Figure S6**). Expressing a tagged protein results in a smaller growth decrease than expressing an untagged protein (**Figure S7**).

## Acknowledgements

We thank Irem Avcilar-Kucukgoze for creating the original GFP-expressing plasmid, Robert Sauer for providing us with the pBAD33-mf-lon plasmid, and Robert Gennis and Toshio Iwasaki for providing us with the ML17 and RF10 *E. coli* strains. This work was supported by European Union’s Horizon 2020 research and innovation program as part of the SynCrop ETN under the Marie Skłodowska-Curie grant 764591 (Z.I.), BrisSynBio, a BBSRC/EPSRC Synthetic Biology Research Centre grant BB/L01386X/1 (T.E.G.), a Royal Society University Research Fellowship grant UF160357 (T.E.G.), and a Turing Fellowship from The Alan Turing Institute under the EPSRC grant EP/N510129/1 (T.E.G.)

## Author contributions

Z.I. and T.E.G. conceived the study. K.S. performed all experiments and analyses. T.E.G. developed the mathematical model. Z.I. and T.E.G. supervised the work and discussed the data. All authors contributed to the writing of the manuscript.

