## Supplementary Information for "Improving the robustness of engineered bacteria to nutrient stress using programmed proteolysis"

**Supporting Figures** **Page**

Figure S1: The effects of Ec and Mf proteolysis tags on cell fluorescence and growth. S2

Figure S2: Off-target degradation by *E. coli* proteases on the Mf tag S3

Figure S3: The specificity of the Lon protease for the Mf tag S4

Figure S4: *E. coli* and *M. florum* proteolysis systems are effective among different *E. coli* strains S5

Figure S5: Programmed proteolysis allows for better cell growth when hosting burdensome genetic circuits S6

Figure S6: The dynamics of the *M. florum* proteolysis system S7

Figure S7: Expressing a tagged protein results in a smaller growth decrease than expressing an untagged protein S8

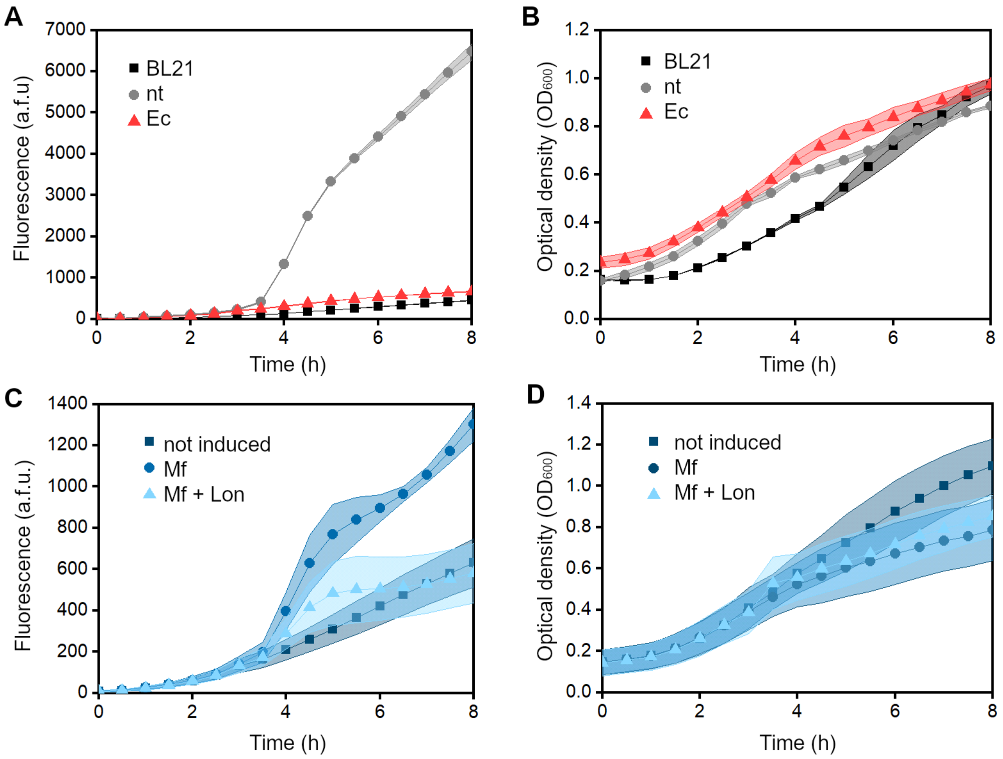


**Figure S1: The effects of Ec and Mf proteolysis tags on cell fluorescence and growth.** (**A**) GFP fluorescence (a.f.u) of untransformed *E. coli* BL21(DE3) cells, or cells transformed with pET-16b-GFP (nt) or pET16b-GFP-Ec (Ec) (**B**) Growth ($OD_{600}$) of untransformed *E. coli* BL21(DE3) cells, or cells transformed with pET-16b-GFP (nt) or pET16b-GFP-Ec (Ec) (**C**) GFP fluorescence (a.f.u) of *E. coli* BL21(DE3) cells co-transformed with pET16b-GFP-Mf and pSB3C5-Mf-Lon (**D**) Growth ($OD_{600}$) of *E. coli* BL21(DE3) cells co-transformed with pET16b-GFP-Mf and pSB3C5-Mf-Lon. Cells were induced with only 0.5 mM IPTG or 0.5mM IPTG and 0.2% (w/v) arabinose (Mf + Lon) at 3 h. Data are means ± SD (*n* = 3 independent biological replicates). Values were used to calculate the normalized fluorescence in **Figure 1B**.

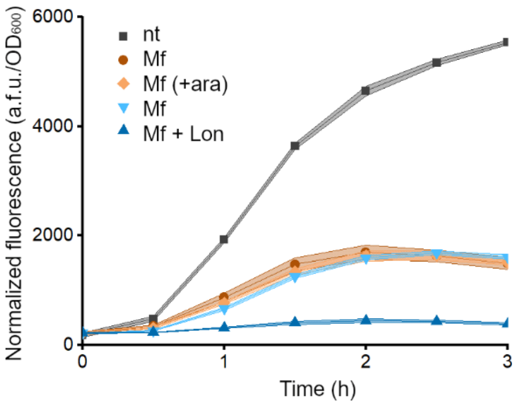


**Figure S2. Off-target degradation by *E. coli* proteases on the Mf tag.** Fluorescence normalized to cell density of *E. coli* BL21(DE3) cells expressing untagged GFP, cells expressing GFP-Mf transformed with only the GFP-Mf expressing plasmid (●,♦), and cells co-transformed with the GFP-Mf and Mf-Lon-expressing plasmids (▼,▲). Cells were induced with 0.5 mM IPTG and 0.2% (w/v) arabinose (Mf + ara, Mf + Lon) at $t$ = 0. Data are means ± SD ($n$ = 3 independent biological replicates).

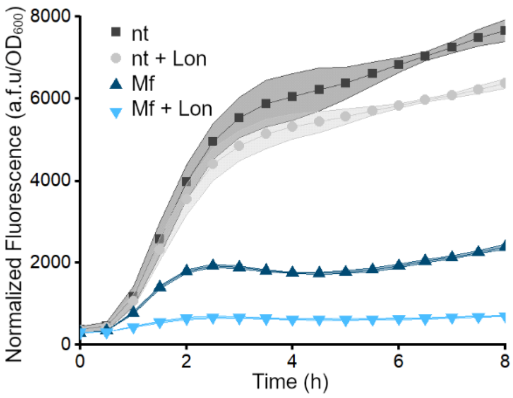


**Figure S3: The specificity of the Lon protease for the Mf tag**. Fluorescence normalized to cell density of *E. coli* BL21(DE3) cells co-transformed with untagged GFP and Mf-Lon -expressing plasmids (■, ●) or GFP-Mf and Mf-Lon -expressing plasmids (▲,▼), and induced to express untagged GFP (nt) or GFP with Mf tag (Mf), with or without co-expressing the Mf-Lon protease (Lon). Cells were induced with 0.5 mM IPTG or 0.5 mM IPTG and 0.2% (w/v) arabinose at *t* = 0. Data are means ± SD (*n* = 3 independent biological replicates).

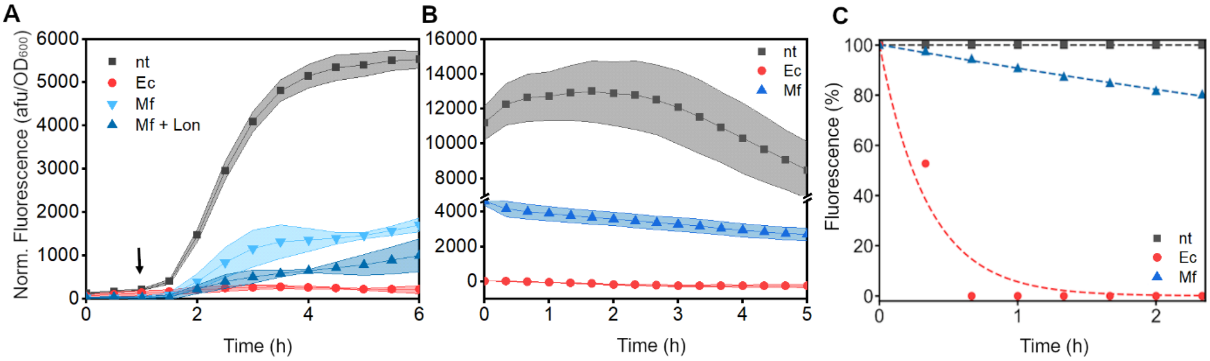


**Figure S4: *E. coli* and *M. florum* proteolysis systems are effective among different *E. coli* strains.** (**A**) GFP fluorescence normalized to cell density of *E. coli* BL21(DE3) star cells expressing non-tagged GFP (nt), GFP-Ec (Ec) or GFP-Mf without and with the co-expression of Mf-Lon (Mf and Mf + Lon, respectively). Arrow indicates timepoint of GFP induction. (**B**) GFP fluorescence normalized to cell density of cells expressing untagged GFP (nt), GFP-Ec (Ec), or GFP-Mf (Mf) after removal of inducer, whilst maintaining Mf-Lon expression in the case of GFP-Mf. (**C**) Percentage fluorescence normalized to the time of removal of the inducer of cells expressing untagged GFP (nt), GFP-Ec (Ec), or GFP-Mf (Mf). Curves are fitted to first order exponential decay. Data are means ± SD (*n* = 3 independent biological replicates).

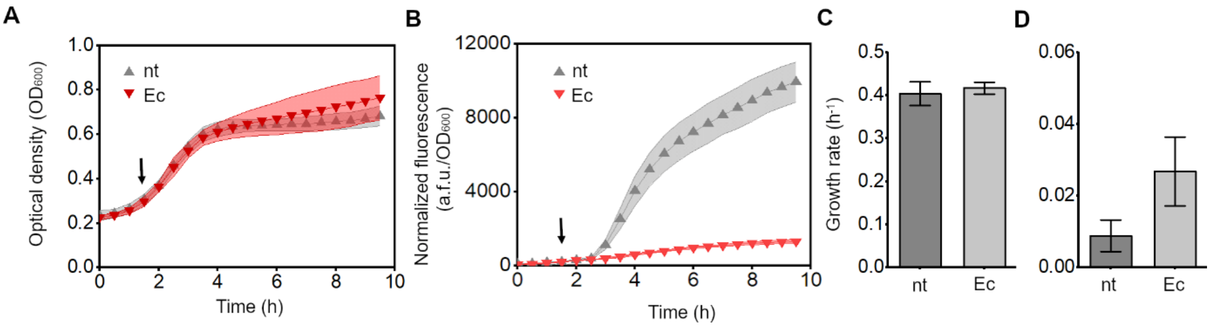


**Figure S5: Programmed proteolysis allows for better cell growth when hosting burdensome genetic circuits.** (**A**) Growth of *E. coli* BL21(DE3) cells induced to express untagged GFP (nt) or GFP-Ec, alongside induction of the 0xF6 plasmid with 2ng/ml aTc. (**B**) Fluorescence of cells induced to express untagged GFP (nt) or GFP-Ec, alongside induction of the 0xF6 plasmid. Arrows indicate timepoint of induction. (**C**) Quantification of the exponential growth rates of cells (1.5–2.5 h). (**D**) Change in cell density ($OD_{600}$) during the slow growth phase (4–9.5 h). Data are means ± SE (*n* = 5 independent biological replicates).

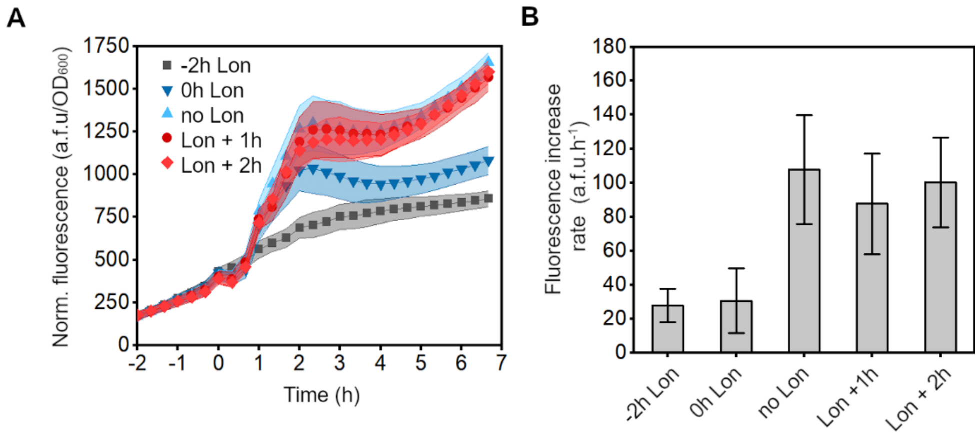


**Figure S6: The dynamics of the *M. florum* proteolysis system.** (**A**) GFP fluorescence normalized to cell density of *E. coli* BL21(DE3) cells expressing GFP-Mf without/with Mf-Lon induced at different time points. GFP-Mf expression was induced with 0.5mM IPTG at $t$ = 0. Mf-Lon expression was induced 2 h before, simultaneously, or 1 or 2 h after GFP-Mf induction, with 0.2% (w/v) arabinose. Data are means ± SD (**B**) Fluorescence increase rate of cells expressing GFP-Mf without/with Mf-Lon induced at different time points in relation to GFP-Mf induction. Data are means ± SE ($n$ = 5 independent biological replicates).

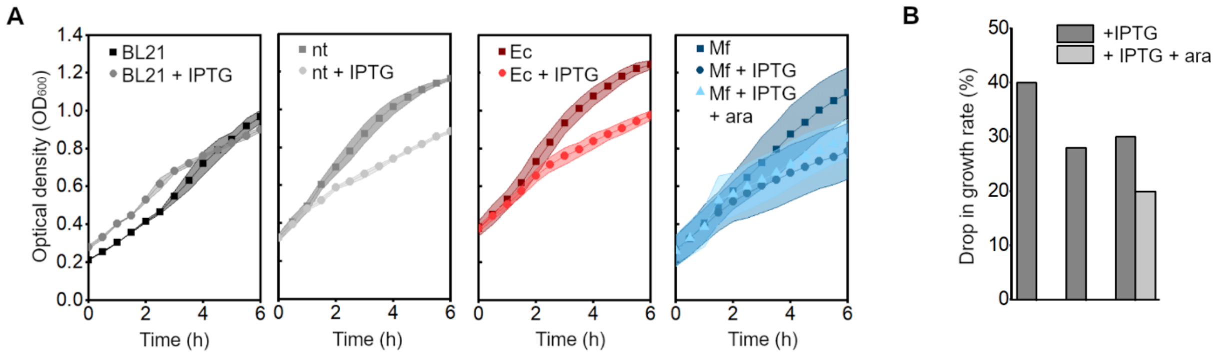


**Figure S7: Expressing a tagged protein results in a smaller growth decrease than expressing an untagged protein.** (**A**) The effect of IPTG inducer (0.5mM) on the growth of untransformed *E. coli* BL21(DE3) cells, cells transformed with pET16b-GFP (nt), cells transformed with pET16b-GFP-Ec (Ec), or the combined effect of IPTG (0.5 mM) and arabinose (0.2% (w/v)) on cells co-transformed with pET16b-GFP-Mf and pSB3C5-Mf-Lon. Cells were induced at 1 h. (**B**) Drop in growth rate (%), between 1 and 4 h, of cells induced to express either GFP-nt, GFP-Ec, GFP-Mf (+ IPTG) or GFP-Mf and Mf-Lon (+ IPTG + ara), compared to uninduced cells. Data are means ± SD (*n* = 3 independent biological replicates).
